# Utility of the SWAN Scale for ADHD Trait-Based Genetic Research: A Validity and Polygenic Risk Study

**DOI:** 10.1101/248484

**Authors:** Christie L. Burton, Leah Wright, Janet Shan, Bowei Xiao, Annie Dupuis, Tara Goodale, S-M Shaheen, Elizabeth C. Corfield, Paul D. Arnold, Russell J. Schachar, Jennifer Crosbie

**Affiliations:** Neurosciences and Mental Health, Hospital for Sick Children, Toronto, Canada; Genetics and Genome Biology Programs, Hospital for Sick Children, Toronto, Canada; Clinical research services; Hospital for Sick Children, Toronto, Canada; Dalla Lana School of Public Health, University of Toronto, Toronto, Canada; Department of Psychiatry, University of Toronto, Toronto, Canada; Institute of Medical Science, University of Toronto, Toronto, Canada; Mathison Centre for Mental Health Research and Education; Departments of Psychiatry and Hotchkiss Brain Institute, Cumming School of Medicine, University of Calgary, Calgary, Canada

**Keywords:** ADHD, SWAN, psychometric validity, polygenic risk score, standardized norms

## Abstract

**Background:** Valid and genetically-informative trait measures of psychopathology collected in the general population would provide a powerful complement to case/control genetic designs. We report the convergent, predictive and discriminant validity of the parent- and the self-report versions of the Strengths and Weaknesses of ADHD Symptoms and Normal Behavior Rating Scale (SWAN) for attention-deficit/hyperactivity disorder (ADHD) traits. We tested if SWAN ADHD scores were associated with ADHD diagnosis, ADHD polygenic risk, as well as with traits and polygenic risk for co-occurring disorders such as anxiety and obsessive-compulsive disorder (OCD).

**Methods:** We collected parent- and self-report SWAN scores in a community sample (n=15,560; 6-18 years of age) and created norms. Sensitivity-specificity analyses determined SWAN cut-points that discriminated those with a community ADHD diagnosis (n=972) from those without a community diagnosis. We validated cut-points from the community sample in a clinical sample (266 ADHD cases; 36 controls). We tested if SWAN scores were associated with anxiety and obsessive-compulsive (OC) traits and polygenic risk for ADHD, OCD and anxiety disorders.

**Results:** Both the parent- and the self-report SWAN measures showed high convergent validity with established ADHD measures and distinguished ADHD participants with high sensitivity and specificity in the community sample. Cut-points established in the community sample discriminated ADHD clinic cases from controls with a sensitivity of 86% and specificity of 94%. High parent- and self-report SWAN scores and scores above the community-based cut-points were associated with polygenic risk for ADHD. High ADHD traits were associated with high anxiety traits, but not OC traits. SWAN scores were not associated with OCD or anxiety disorder polygenic risk.

**Conclusion:** The parent- and self-report SWAN are potentially useful in genetic research because they predict ADHD diagnoses and are associated with ADHD polygenic risk.

---

Attention-deficit/hyperactivity disorder (ADHD) is an impairing and common disorder (5% prevalence; Polanczyk, de Lima, Horta, Biederman, & Rohde, 2007). Despite high heritability (70-80%; Faraone et al., 2015), few genetic variants have been identified (Demontis et al., in press). An obstacle to more rapid progress is the high cost and slow speed of collecting sufficiently powered clinical samples. Most genome-wide association studies (GWAS) compare controls to cases, identified by clinical interviews using diagnostic criteria. This approach does not consider phenotypic variation among cases or controls. Some cases are more severe than others, while controls can range from average to above average in their attention, impulse control and activity level (Fair, Bathula, Nikolas, & Nigg, 2012). Most symptom-based scales, which typically generate skewed distributions, are often used as quantitative trait measures or are dichotomized into case/control. Both approaches can reduce statistical power (van der Sluis, Posthuma, Nivard, Verhage, & Dolan, 2013). Skewed symptom counts are more likely and problematic in the general population because of the low base-rate of most disorders, which results in a clustering of scores at zero. To accelerate genetic discovery in ADHD, we need genetically-informative phenotype measures that efficiently generate reliable, heritable, widely and preferably normally distributed scores in the general population where samples can be more rapidly accrued.

The Strengths and Weaknesses of ADHD Symptoms and Normal Behavior Rating Scale (SWAN; Swanson et al., 2012) is a unique ADHD trait measure. The SWAN is based on the ADHD criteria from the Diagnostic and Statistical Manual of Mental Disorders, 4^th^ edition (American Psychiatric Association, 2013). What makes the SWAN different is that it rates symptoms on a seven-point Likert scale with −3 indicating low ADHD traits and +3 indicating high ADHD traits. The SWAN meets many criteria of a genetically-informative trait measure. It is reliable, generates normally distributed scores in the general population, has high internal consistency, high test-retest reliability, converges reasonably with ADHD symptom measures and diverges from measures of emotionality (Arnett et al., 2013; Crosbie et al., 2013; Lai et al., 2013; Lakes, Swanson, & Riggs, 2012; Stroud et al., 2009; Swanson et al., 2012). Twin and sibling studies established the heritability (h^2^) of the parent- and self-report SWAN (h^2^=0.24-0.94; Couvy-Duchesne et al., 2016; Crosbie et al., 2013; Greven et al., 2016; Hay, Bennett, Levy, Sergeant, & Swanson, 2007; Peng et al., 2016; Polderman et al., 2007; Smit & Anokhin, 2017). A GWAS using the SWAN identified suggestive genome-wide significant hits (n=1851; Ebejer et al., 2013).

Questions remain about the SWAN’s suitability for genetic research. First, it is unclear whether the SWAN measured in the general population can discriminate ADHD cases from controls with adequate sensitivity and specificity. The question is not whether SWAN can substitute for a diagnosis of ADHD, but whether the extreme of widely distributed ADHD traits converge with ADHD diagnosis (Crosbie et al., 2013; Plomin, Haworth, & Davis, 2009). People with extremely high traits should have ADHD, while people with low trait scores should not have ADHD. Second, it is unclear if low ADHD traits, such as hyper-attentiveness and lack of fidgeting, may represent a strength or could represent another disorder. Anxiety or obsessive-compulsive disorder (OCD) are partially characterized by over-attention to threatening stimuli or underactivity due to shyness or fear (Muller & Roberts, 2005; Roy et al., 2008). Third, a validated self-report version of the SWAN for use in youth is critical to optimize utility in population genetics research given that parents are not always available. While a self-report version has been used previously in youth (Couvy-Duchesne et al., 2016; Ebejer et al., 2015; Greven et al., 2016), it was not previously validated. Finally, we do not know if parent- and self-report SWAN scores are sensitive to polygenic risk for ADHD (i.e., ADHD polygenic risk increases with SWAN scores) and not polygenic risk for co-occurring psychopathology such as OCD or anxiety disorders (Abramovitch, Dar, Mittelman, & Wilhelm, 2015; Jensen et al., 2001).

The current study expands on existing psychometrics of the SWAN with a focus on its potential utility for genetic research. We examined the convergent, predictive and discriminant validity in parent- and self-report versions of the scale, extracted norms and clinical cut-points. We tested if the SWAN discriminated youth with, compared to without, an ADHD diagnosis in a large community sample and in a clinic sample. We also validated a self-report version of the SWAN. We tested the hypothesis that high ADHD traits would be associated with elevated anxiety and OC traits based on their co-occurrence with ADHD (Abramovitch et al., 2015; Jensen et al., 2001) as well as whether anxiety traits and obsessive-compulsive (OC) traits were associated with low ADHD trait scores. We also examined if participants with high SWAN scores had higher ADHD, but not OCD or anxiety disorder, polygenic risk scores.

## Methods

### Community sample

We recruited 17,263 participants (ages 6-18 years) at the Ontario Science Centre in Toronto, Canada (see Crosbie et al., 2013 for details). Complete information about demographics, history of community diagnosis of ADHD and ADHD traits was available for 15,560 participants (parent-report: ages 6-16 years n=12,797, self-report: ages 13-18 years n=2,763). Consistent with the reported prevalence of childhood ADHD (CDC, 2015), 972 (6.2%) reported a diagnosis of ADHD (referred to as community diagnosis).

The parent-report SWAN scale (SWAN-Parent) consisted of 18 items with a hyperactive/impulsive (SWAN-Parent-HI) and inattentive subscale (SWAN-Parent-IA) as well as a combined total (SWAN-Parent-Com). The self-report version of the SWAN scale (SWAN-Self) changed the referent to “me” rather than “your child”. SWAN-Self consisted of the same two subscales (SWAN-Self-HI, SWAN-Self-IA) and a combined total score (SWAN-Self-Com).

To assess convergent validity, a random subset of participants completed a widely-used ADHD rating scale: Conners’ ADHD Rating Scale-Revised (CPRS-R; n=841) or Conners-Wells Adolescent Self Report Scale (CASS-L; n=172). Each measure generates four scales in T-score format: The Inattentive scale (L-scale), Hyperactive-Impulsive scale (M-scale), Total scale (N-scale) and an ADHD Index scale (H-scale) (Conners, Sitarenios, Parker, & Epstein, 1998).

Parents and youth completed questionnaires that measure anxiety (Child Behavior Checklist [CBCL] anxiety problems sub-scale; Kendall et al., 2007) and OC traits (Toronto Obsessive-Compulsive Scale – TOCS; Park et al., 2016). The anxiety scale has 11 items ranging from 0 (“not true”) to 2 (“very true or often true”). The TOCS has 21-items scored on a seven-point Likert scale ranging from −3 (“far less often than others of the same age”) to +3 (“far more often than others of the same age”).

### ADHD Clinic Sample

The clinic sample consisted of 266 children (6-18 years old) with a diagnosis of ADHD and 36 control children assessed in a tertiary care mental health clinic. ADHD diagnoses were based on consensus between a psychiatrist and clinical psychologist following a rigorous assessment described elsewhere (McAuley, Chen, Goos, Schachar, & Crosbie, 2010). We excluded individuals with an IQ <80 on both verbal and non-verbal domains. SWAN-Parent scores were not considered as part of the diagnosis and SWAN-Self were not collected because most participants were pre-adolescent.

### Ethical Considerations

Informed consent, and verbal assent when applicable, approved by The Hospital for Sick Children Research Ethics Board were obtained from all participants.

### Statistical analyses

The SWAN items were reversed such that high scores reflected high ADHD traits and low scores reflected low ADHD traits. We standardized SWAN scores for age and gender to account for their well-established association with ADHD traits (Ramtekkar, Reiersen, Todorov, & Todd, 2010; details in supplemental material). Standardized scores were created for the total score (zSWAN-Com) and each subscale (zSWAN-IA and zSWAN-HI), TOCS total score and CBCL anxiety problem total scores. Bootstrap analysis using SAS 9.3 established confidence intervals. We accounted for sibling relatedness in the models using random effects.

We used t-tests to compare SWAN scores in participants with and without a community-reported diagnosis of ADHD. In analyses without standardized zSWAN scores, we included age and gender as covariates in parent- and self-respondents separately.

Receiver operating characteristic curves (ROC) selected optimal models for discriminating those with a community ADHD diagnosis. Area under the curve (AUC) of ≥0.80 indicates good discrimination of cases from controls. The Youden Index indicates the optimal cut-points in an ROC curve. We validated the ability of cut-points from the community sample to correctly classify ADHD cases in the clinic sample. We compared the predictive validity of community-derived cut-points with cut-points recommended by Swanson et al. (1.65 SD; 2012). ROC analyses were conducted using MedCalc Application. All other statistical tests were performed using SPSS 21 and SAS v9.4.

Internal consistency was assessed using Cronbach’s α for both the SWAN-Parent and SWAN-Self (α ≥0.80 are considered good). To assess convergent validity, z-SWAN subscales were correlated with their corresponding CPRS-R and CASS-L subscales in 841 participants who completed two measures. Spearman’s rho was used to assess correlations because CPRS-R and CASS-L scores were not normally distributed.

To examine the relationship of ADHD traits with OC and anxiety traits, we divided participants into five groups based on zSWAN-Com scores (group 1 [low traits]: n=357, group 2: n=3,607, group 3: n=7,739, group 4: n=3,515 and group 5 [high traits]: n=342). Analyses of Variance (ANOVAs) were used to assess mean differences in standardized TOCS total score and CBCL anxiety total score across SWAN groups.

We genotyped a subset of the sample (n=5,366; details in online supplement) and used 5,154 participants that passed standard quality control in the analyses. Polygenic risk scores were calculated based on three discovery samples: ADHD from the Psychiatric Genomics Consortium meta-analysis (cases=20,183, controls=35,191; Demontis et al., in press), OCD from the International OCD Foundation Genetics Collaborative and OCD Collaborative Genetics Association Studies meta-analysis (cases=2688 and controls=7037; International Obsessive Compulsive Disorder Foundation Genetics Collaborative & Studies, 2017) and anxiety disorders from the Anxiety Neuro Genetics Study (ANGST; 17,310 cases and controls; Otowa et al., 2016). From each discovery set, we selected a subset of pruned SNPs based on a range of p-value thresholds (p<1×10^−5^, 1×10^−4^, 1×10^−3^, 0.05, 0.01, 0.10, 0.20, 0.30, 0.40 and 0.50). Pruning was conducted in plink 1.9 (Purcell et al., 2007; http://pngu.mgh.harvard.edu/purcell/plink/ref) on unambiguous variants from the discovery sets. The standardized polygenic risk scores (mean=0, SD=1) were sums of the allele counts from our community-based sample weighted by the effect size from each discovery set.

We developed a script based on PRSice (Euesden, Lewis, & O′Reilly, 2015) to test the association between zSWAN-Com scores with polygenic risk scores at each p-value threshold while correcting for potential effects of array and population stratification using principal components (see supplemental methods). We selected the p-value threshold that accounted for the most variance in zSWAN using r^2^ for each discovery set for subsequent analyses. To understand if polygenic risk for ADHD, anxiety and OCD were associated with high and low SWAN scores, we divided genotyped participants into three equal groups based on their zSWAN-Com scores (low, medium, high) and compared the pair-wise mean polygenic risk score differences across the groups using Tukey’s test within each discovery set (ADHD, OCD and anxiety). We also used a two-sided t-test to compare polygenic risk scores derived from the ADHD discovery set in participants above and below the optimal cut-points identified in the ROC analyses for zSWAN-Com for parent-report (n=4,426) and self-report data (n=728) separately. We conducted similar analyses using the Swanson recommended cut-point (1.65 SD; Swanson et al., 2012) in all genotyped participants (n=5,154). The significance alpha threshold was adjusted to account for multiple testing (0.05/41=0.001).

## Results

SWAN-Parent and SWAN-Self scores were significantly higher in participants with, compared to without, an ADHD community diagnosis (Table 1). Community and clinic ADHD cases had comparable SWAN scores while clinic controls had lower SWAN scores than community controls (Table 1 and 2). SWAN scores showed high sensitivity and specificity for community-reported ADHD diagnosis (Table 3). Sensitivity and specificity were good for both scales but greater for SWAN-Parent (AUC=0.85-0.90) than SWAN-Self (AUC=0.71-0.76). Differences between zSWAN-Com and non-standardized SWAN scores in predicting ADHD community diagnoses were small. Sensitivity and specificity were reduced when we applied the cut-point suggested by Swanson et al. (2012) for zSWAN–Com and SWAN-Parent-Com in the community sample (sensitivity 57% and 70% respectively; specificity 92% and 90% respectively).

**Table 1.** Mean Parent- and Self-report SWAN by reported ADHD diagnosis (Community sample)

|  | Without community<br>ADHD diagnosis | With community<br>ADHD diagnosis | *t*-value | Cohen's *d* |
| --- | --- | --- | --- | --- |
| Total N | 14,588 | 972 |  |  |
| **Parent-report SWAN** |  |  |  |  |
| N | 11,987 | 810 |  |  |
| Age (SD) | 10.06 (2.08) | 10.77 (2.08) |  |  |
| Male (N, %) | 6203 (51.75) | 613 (75.68) |  |  |
| **SWAN-Parent** |  |  |  |  |
| SWAN-Parent-Com (SD) | -6.55 (16.06) | 20.29 (14.52) | -45.21* | 1.68 |
| SWAN-Parent-IA (SD) | -2.64 (8.66) | 10.99 (8.20) | -41.87* | 1.58 |
| SWAN-Parent-HI (SD) | -3.91(8.89) | 9.30 (8.59) | -40.40* | 1.49 |
| **zSWAN** |  |  |  |  |
| zSWAN-Com (SD) | -0.09 (0.94) | 1.37 (0.79) | -43.41* | 1.57 |
| zSWAN-IA (SD) | -0.09 (0.95) | 1.27 (0.84) | -39.74* | 1.44 |
| zSWAN-HI (SD) | -0.08 (0.95) | 1.25 (0.85) | -39.37* | 1.36 |
| **Self-report SWAN** |  |  |  |  |
| N | 2,601 | 162 |  |  |
| Age (SD) | 15.34 (1.29) | 15.41(1.22) |  |  |
| Male (N, %) | 983 (37.79) | 104 (64.20) |  |  |
| **SWAN-Self** |  |  |  |  |
| SWAN-Self-Com (SD) | -5.58 (13.96) | 8.06 (14.27) | -12.26* | 0.98 |
| SWAN-Self-IA (SD) | -3.00 (7.55) | 3.85 (7.69) | -11.37* | 0.91 |
| SWAN-Self-HI (SD) | -2.58 (8.47) | 4.21 (8.86) | -10.00* | 0.80 |
| **zSWAN** |  |  |  |  |
| zSWAN-Com (SD) | -0.05 (0.97) | 0.88 (0.98) | -11.87* | 0.96 |
| zSWAN-IA (SD) | -0.05 (0.98) | 0.80 (0.97) | -10.70* | 0.87 |
| zSWAN-HI (SD) | -0.05 (0.98) | 0.75 (1.04) | -9.99* | 0.81 |

\* $p < 0.0001$ . SWAN = Strengths and Weaknesses of ADHD Symptoms and Normal Behavior
Rating Scale, SD = standard deviation; SWAN-Parent-Com = Parent-report SWAN, combined;
SWAN-Parent-IA = Parent-report SWAN, inattentive subscale; SWAN-Parent-HI = Parent-
report SWAN, hyperactive/impulsive subscale; SWAN-Self-Com = Self-report SWAN,
combined; SWAN-Self-IA = Self-report SWAN, inattentive subscale; SWAN-Self-HI = Self-
report SWAN, hyperactive subscale; zSWAN-Com = Standardized SWAN score, combined;
zSWAN- IA = Standardized SWAN for both informants, inattentive subscale; zSWAN-HI = Standardized SWAN, hyperactive/impulsive subscale.

**Table 2.** Mean parent-report SWAN scores in ADHD cases and controls (Clinic sample)

|  | **Controls** | **ADHD cases** | ***t*-value** | **Cohen's *d*** |
| --- | --- | --- | --- | --- |
| N | 36 | 266 |  |  |
| Age (SD) | 8.92 (1.87) | 9.15 (2.26) |  |  |
| Male (N, %) | 16 (44.44) | 209 (78.57) |  |  |
| **<u>SWAN-Parent</u>** |  |  |  |  |
| SWAN-Parent-Com (SD) | -14.47 (17.98) | 20.35 (13.27) | -11.21* | 2.50 |
| SWAN-Parent-IA (SD) | -7.44 (9.85) | 12.19 (8.26) | -13.07* | 2.32 |
| SWAN-Parent-HI (SD) | -7.03 (9.13) | 8.16 (8.16) | -10.33* | 1.83 |
| **<u>zSWAN</u>** |  |  |  |  |
| zSWAN-Com (SD) | -0.51 (1.05) | 1.36 (0.68) | -10.38* | 2.55 |
| zSWAN-IA (SD) | -0.55 (1.08) | 1.38 (0.83) | -10.35* | 2.24 |
| zSWAN-HI (SD) | -0.41 (1.00) | 1.12 (0.78) | -8.80* | 1.89 |

\* $p < 0.0001$ ; no self-report scales were collected in this sample; SWAN = Strengths and Weaknesses of ADHD Symptoms and Normal Behavior Rating Scale, SD = standard deviation; SWAN-Parent-Com = Parent-report SWAN, combined; SWAN-Parent-IA = Parent-report SWAN, inattentive subscale; SWAN-Parent-HI = Parent-report SWAN, hyperactive/impulsive subscale; zSWAN-Com = Standardized SWAN score, combined; zSWAN- IA = Standardized SWAN for both informants, inattentive subscale; zSWAN-HI = Standardized SWAN, hyperactive/impulsive subscale.

**Table 3.** Area under the curve (AUC), optimal cut-points, sensitivity and specificity for classifying community-diagnosed ADHD (Community sample)

|  | AUC | Optimal cut-point | Sensitivity | Specificity |
| --- | --- | --- | --- | --- |
| **Parent-report** |  |  |  |  |
| SWAN-Parent-Com | 0.90 | >6 | 83.7 | 83.3 |
| SWAN-Parent-IA | 0.88 | >4 | 78.6 | 82.9 |
| SWAN-Parent-HI | 0.86 | >2 | 77.2 | 81.0 |
| zSWAN- Com | 0.88 | >0.74 | 82.5 | 81.0 |
| zSWAN- IA | 0.86 | >0.60 | 81.2 | 76.3 |
| zSWAN- HI | 0.85 | >0.72 | 77.0 | 81.0 |
| **Self-report** |  |  |  |  |
| SWAN-Self-Com | 0.76 | >4 | 59.9 | 78.9 |
| SWAN-Self-IA | 0.74 | >0 | 66.7 | 69.0 |
| SWAN-Self-HI | 0.71 | >2 | 58.6 | 75.2 |
| zSWAN-Com | 0.75 | >0.81 | 57.4 | 81.4 |
| zSWAN-IA | 0.73 | >0.51 | 64.2 | 71.4 |
| zSWAN-HI | 0.71 | >0.56 | 61.7 | 73.1 |

SWAN-Parent-Com = Parent-report SWAN, combined; SWAN-Parent-IA = Parent-report SWAN, inattentive subscale; SWAN-Parent-HI = Parent-report SWAN, hyperactive/impulsive subscale; SWAN-Self-Com = Self-report SWAN, combined; SWAN-Self-IA = Self-report SWAN, inattentive subscale; SWAN-SELF-HI = Self-report SWAN, hyperactive subscale; zSWAN-Com = Standardized SWAN score for both informants, combined; zSWAN- IA = Standardized SWAN for both informants, inattentive subscale; zSWAN-HI = Standardized SWAN for both informants, hyperactive/impulsive subscale

In the clinic sample, a SWAN-Parent-Com cut-point of >6 resulted in sensitivity of 86% and specificity of 94%, correctly identifying 228 of 266 ADHD cases (86%) while misclassifying one control into the ADHD group (3%).

SWAN-Parent and SWAN-Self scores showed high internal consistency (SWAN-Parent-Com α=0.95, SWAN-Parent-IA α=0.92 and SWAN-Parent-HI α=0.93; SWAN-Self-C α=0.88, SWAN-Self-IA α=0.82 and SWAN-Self-HI α=0.84). For SWAN-parent, convergence was high between corresponding zSWAN-Com and CPRS-R scale scores: CPRS-R Inattentive and zSWAN-IA (rho=0.70, *p*<0.01), CPRS-R Total and zSWAN-Com (rho=0.72, *p*<0.01), CPRS-R ADHD Index and zSWAN-Com (rho=0.71, *p*<0.01). CPRS-R Hyperactive/Impulsive scale had a slightly lower convergence with the zSWAN-HI (rho=0.67, *p*<0.01). For SWAN-self, convergence was moderate for the zSWAN-IA and the CASS-L Inattention subscale (rho=0.52, *p*<0.01) and the zSWAN-HI and the CASS-L Hyperactivity/Impulsivity subscale (rho=0.58, *p*<0.01).

There was a significant, non-linear association of SWAN scores with TOCS scores (*F(4,6)*=8.45, *p*<.0001; Figure S1a, online supplement), although the effect size was small (Cohen’s d=0.33). Higher zSWAN-Com scores were associated with higher CBCL anxiety total scores (*F(4,6)*=18.33, *p*<.0001). Anxiety traits were significantly higher in the group with the highest z-SWAN-Com scores compared to the lowest (group 1 vs. 5: *p*<0.01; Cohen’s d=0.85) or intermediate groups (group 3 vs. 5: *p*<0.01; Cohen’s d<0.67; Figure S1b, online supplement).

Figure 1 shows that ADHD polygenic risk was significantly associated with zSWAN-Com scores and the p-value = 0.30 threshold from the ADHD discovery set explained the most variance (r^2^=8.74×10^−3^ *p*=1.73×10^−11^). ADHD polygenic risk significantly predicted zSWAN-Com scores in both parent- and self-report groups (*p*=0.00057 and *p*<0.00001 respectively, data not shown). Neither polygenic risk scores based on OCD nor anxiety disorders predicted zSWAN-Com. The p-value thresholds that explained the most variance in each trait were 0.01 for OCD (r^2^=2.27×10^−4^) and 10^−5^ for anxiety (r^2^=1.98×10^−4^). The p-value threshold that explained the most variance for each discovery set was used in the subsequent analyses. ADHD polygenic risk was significantly higher in participants with the highest SWAN scores than in those with low and mid-range scores (low vs. high, *p*=1.48×10^−10^; medium vs. high, *p*=4.27×10^−5^; medium vs. low, *p*=0.07; Figure 2). There were no significant differences in anxiety or OCD polygenic risk scores across the SWAN categories respectively (*p*>0.40, Figure S2, online supplement).

**Figure 1:**
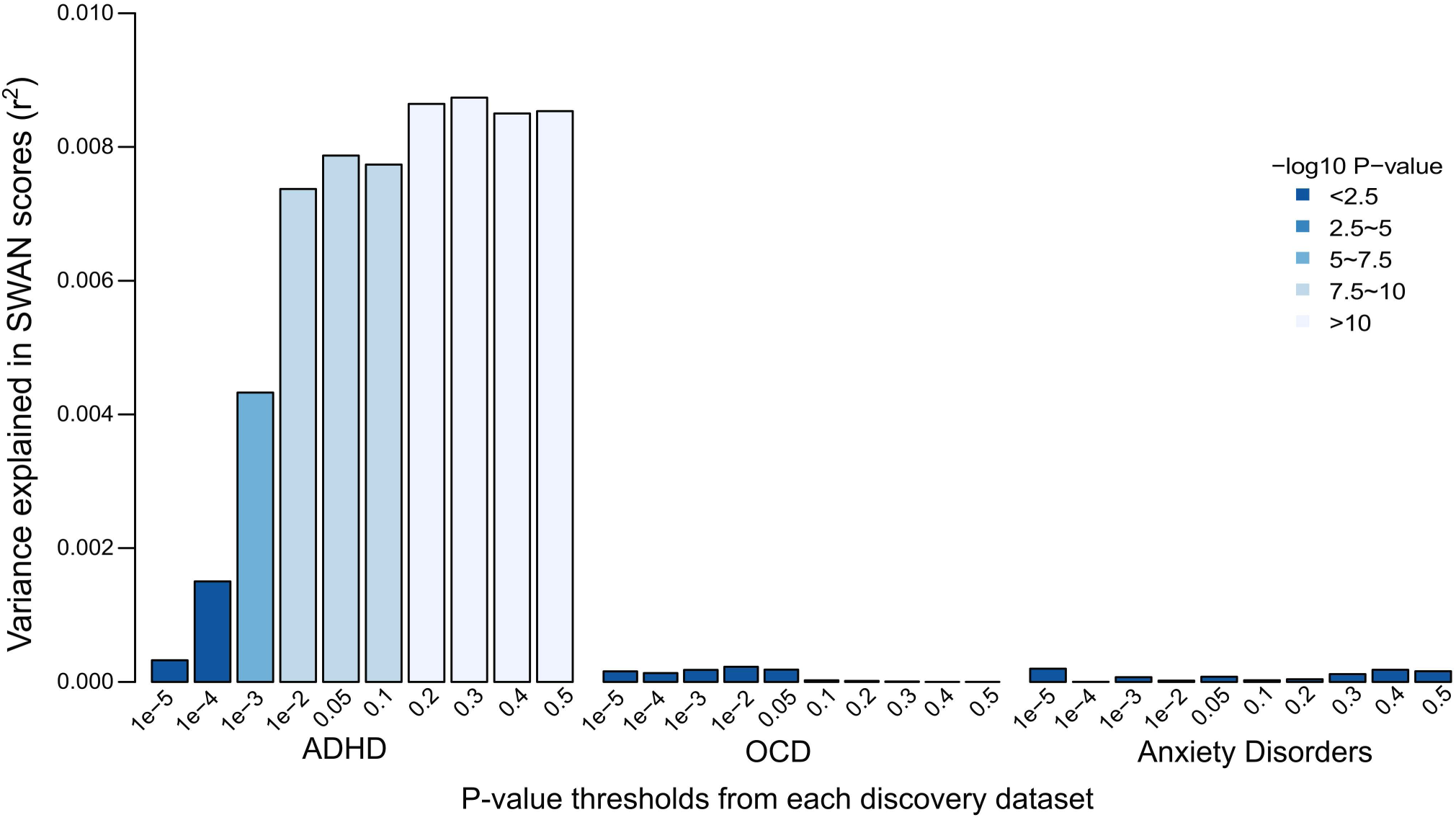
Predicting SWAN Scores from Polygenic Risk for ADHD, OCD and Anxiety. Polygenic risk scores derived from clinical attention-deficit/hyperactivity disorder (ADHD), but not obsessive-compulsive disorder (OCD) or anxiety disorder, discovery samples were significantly associated with ADHD traits (standardized total Strengths and Weaknesses of ADHD Symptoms and Normal Behavior Rating Scale combined [zSWAN-Com] score). P-value thresholds refer to parameters from discovery sample statistics (ADHD, OCD, Anxiety disorders). r^2^ = variance explained by polygenic risk in predicting zSWAN-Com. –log10 p-value reflects the association of polygenic risk scores and zSWAN-Com.

**Figure 2.**
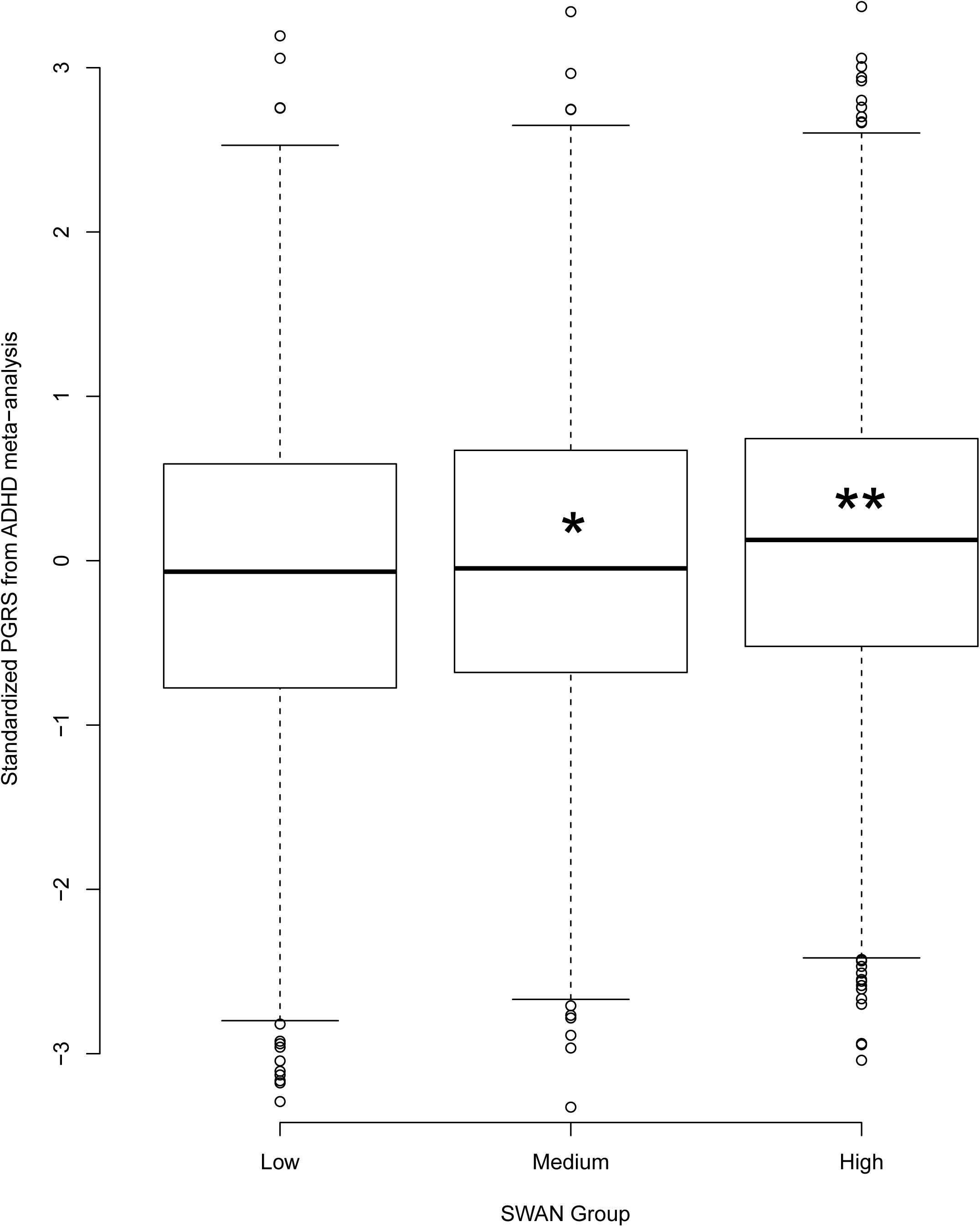
ADHD Polygenic Risk Scores across Low to High ADHD traits. Attention-deficit/hyperactivity disorder (ADHD) polygenic risk was highest in the group with the highest standardized Strengths and Weaknesses of ADHD Symptoms and Normal Behavior Rating Scale combined (zSWAN-com) scores (** low vs. high group *p*=1.48×10^−10^; * medium vs. high, *p*=4.27×10^−5^; medium vs. low, *ns*). n=1,805 in each SWAN group. PGRS = polygenic risk score.

ADHD polygenic risk was significantly higher in participants above, compared to below, the optimal cut-points identified in the ROC analyses for parent-report zSWAN (score≥0.74: t=-5.4; *p*=8.8×10^−8^) and all participants with the Swanson cut-point (using zSWAN-Com score≥1.65: t=3.5; p<0.001; data not shown). The same trend was observed for the cut-point for self-report zSWAN (score≥0.81, t=-1.8; *p*=0.07; data not shown).

## Discussion

Ascertaining sufficiently large samples for well-powered GWAS analyses is expensive and time-consuming. Alternative strategies that use valid and genetically-informative quantitative trait measures to collect population-based samples could accelerate genetic discovery and manage costs. The SWAN appears to be suitable for this purpose based on current and previous results. The SWAN generates heritable trait scores that are widely and virtually normally distributed in the general population (Couvy-Duchesne et al., 2016; Crosbie et al., 2013; Greven et al., 2016; Hay et al., 2007; Peng et al., 2016; Polderman et al., 2007; Smit & Anokhin, 2017). To add support to the validity of the SWAN, we showed that high SWAN scores converge with a clinical diagnosis of ADHD consistent with the quantitative trait notion of psychopathology and SWAN (parent- and self-report) scores show convergent and divergent validity against other measures of ADHD, anxiety and OCD traits. We demonstrated that low ADHD trait scores were not associated with either OCD or anxiety traits. Finally, we found that both parent- and self-report SWAN scores were associated with ADHD, but not OCD or anxiety disorder, polygenic risk.

The current study uses the largest sample to date to confirm the high internal consistency of the SWAN as previously reported (Lai et al., 2013; Lakes et al., 2012; Stroud et al., 2009). We found high convergent validity of the SWAN with a gold-standard measure of ADHD and divergent validity from measures of anxiety and OCD. SWAN scores, whether parent- or self-report, discriminated those with and without a community diagnosis of ADHD with high sensitivity and specificity supporting the validity of the SWAN as a measure of ADHD traits. The optimal cut-point derived in the community sample discriminated ADHD cases from controls in a clinic sample. Prediction of an ADHD diagnosis from SWAN scores was imperfect, which is not surprising given the range of information and informants required for a diagnosis. The cut-points for the SWAN-Parent (score >6) derived from the community sample predicted diagnosis better than the cut-point recommended by Swanson et al. (1.65 SD; 2012). The Swanson cut-point is higher than our identified cut-point resulting in higher specificity, but markedly lower sensitivity. The SWAN-Self in adolescents also had good sensitivity and specificity, high internal consistency and good convergence with an accepted self-report ADHD scale – CASS-L. Finally, both parent- and self-report SWAN scores were associated with ADHD polygenic risk. However, parent- and self-report ADHD traits in youth may have different genetic architectures. The non-significant increase in ADHD polygenic risk in self-report participants above the ROC-derived cut-point was likely related to power because the group was much smaller (>0.81 n=154, <0.81 n=574). Together our results suggest that although correlations between parent- and self-report ADHD symptoms are often low to moderate (Parker, Bond, Reker, & Wood, 2005), the SWAN self-report is a valid measure of ADHD traits and is associated with ADHD genetic risk.

SWAN scores predicted polygenic risk for ADHD (Demontis et al., in press), which supports the hypothesis that ADHD traits measured by the SWAN share genetic risks with ADHD clinical diagnoses. This adds to the mounting evidence that ADHD traits and diagnosis share genetic risk (Groen-Blokhuis et al., 2014; Stergiakouli et al., 2015). As in all other polygenic risk studies, the variance explained in the trait were small, which is likely the result of several factors including the size of discovery and target sample sets and the role of rare variants, epistasis and gene by environment interactions (Mistry, Harrison, Smith, Escott-Price, & Zammit, 2018; van der Sluis et al., 2013).

One critique of the use of trait measures such as the SWAN in research or clinical practice is the presumption that the low extreme of a trait represents a strength (Plomin et al., 2009). Low ADHD traits could reflect above average impulse, motor and attention control (Fair et al., 2012) rather than hypo-activity, inertia, over-focusing, or perseveration seen in OCD or anxiety disorders. In our study, participants with low trait ADHD scores did not have elevated scores for anxiety or for OC traits, supporting the hypothesis that low ADHD trait scores are most likely strengths rather than evidence of a different disorder. The finding that SWAN scores were not associated with polygenic risk for anxiety or OCD further supports this hypothesis.

Trait-based questionnaires can play an important role in clinical practice and research. Questionnaires with appropriate age and gender norms could be useful for screening of ADHD symptoms as part of a comprehensive assessment, for establishing a treatment baseline and for monitoring progress. However, a diagnosis of ADHD requires a comprehensive clinical assessment and should not be based solely on questionnaire information, especially from a single informant.

Our study has various strengths. It is the largest population-based study of the SWAN and the first study to calculate general population-based cut-points and validate them in a clinic sample. Age and gender standardized SWAN scores were used to create scoring software, the ADHD Calculator of Traits (ACT©; http://www.sickkids.ca/Research/schachar-lab/index.html).

Limitations of our study include that our sample was slightly biased towards higher socio-economic status (SES); however, SES was not associated with traits in our sample (Crosbie et al., 2013). We were also unable to assess the association of ADHD traits with conduct and oppositional-defiant disorder traits, as they were not measured in our community sample. Future studies should examine genetic overlap between ADHD and these traits. Finally, our discovery samples for the polygenic risk score analyses included youth and adult participants while our target sample included youth only. We are currently unable to divide discovery sets based on age of onset. Age of onset in discovery sets may not have affected our results given that genetic risk for adult disorders like schizophrenia are associated with traits related to childhood disorders (Pain et al., 2018; Riglin et al., 2017) and ADHD is comorbid with OCD and anxiety in adults (Tan, Metin, & Metin, 2016; Yang, Tai, Yang, & Gau, 2013).

## Conclusion

Our study adds support to the validity of the SWAN as a valid quantitative trait measure that is sensitive to genetic variation in ADHD, but not genetic variation in other disorders. Our results suggest the utility of the SWAN to collect ADHD traits in population-based designs to speed genetic discovery and reduce collection costs.

#### Key Points

- Valid and genetically-informative quantitative trait measures in the community are a viable alternative for gene discovery in ADHD
- Both the parent- and self-report SWAN scales (Strengths and Weaknesses of ADHD Symptoms and Normal Behavior Rating Scale) had sound psychometric properties
- High ADHD traits were associated with ADHD while low ADHD traits were not associated with anxiety or OCD
- ADHD traits shared polygenic risk with diagnosed ADHD

## Acknowledgements

This study was funded by the Canadian Institutes of Health Research (PDA: MOP-106573; RJS: MOP–93696) and the Endowment Fund from the Department of Psychiatry, Hospital for Sick Children. We thank the Psychiatric Genomics Consortium ADHD working group, the International Obsessive Compulsive Disorder Foundation Genetics Collaborative and OCD Collaborative Genetics Association Studies and Anxiety Neuro Genetics Study consortium for allowing us to use their summary statistics. We thank Lisa Strug for her statistical consultation and all individuals involved in collecting the community and clinic datasets.

## Abbreviations

OC: Obsessive-compulsive
SWAN: Strengths and Weaknesses of ADHD Symptoms and Normal Behavior Rating Scale
SWAN-Parent: SWAN parent-report scale
SWAN-Parent-HI: SWAN parent-report hyperactive/impulsive sub-scale
SWAN-Parent-IA: SWAN parent-report inattentive subscale
SWAN-Parent-Com: SWAN parent-report combined total score
SWAN-Self: SWAN self-report scale
SWAN-Self-HI: SWAN self-report hyperactive/impulsive sub-scale
SWAN-Self-IA: SWAN self-report inattentive subscale
SWAN-Self-Com: SWAN self-report combined total score
CPRS-R: Conners’ ADHD Rating Scale-Revised
CASS-L: Conners-Wells Adolescent Self Report Scale
CBCL: Child Behavior Checklist
TOCS: Toronto Obsessive-Compulsive Scale
zSWAN-Com: SWAN total z-score
zSWAN-IA: SWAN inattentive subscale z-score
zSWAN-HI: SWAN hyperactive/impulsive z-scores

